# Methodological Impacts on Microbiome Structure and Indicator Status in the Human Lower Respiratory Tract

**DOI:** 10.64898/2026.08.02.742334

**Authors:** Avery J. C. Noonan, Renelle Myers, Ryan McLaughlin, Anika Nag, Steven Q. Chen, Crista Bartolomeu, Scott A. Borden, Stephen Lam, Steven J. Hallam

## Abstract

**Rationale:** Lung cancer is the leading cause of cancer-related death globally, and rising incidence among traditionally low-risk individuals intensifies the need for improved early-detection methods that the lower-airway microbiome may inform.

**Objectives:** To evaluate how respiratory tract sampling method shapes inferred microbiome structure, and whether bronchial brushing recovers a microbial community ecologically distinct from BAL and oral rinse.

**Methods:** A pilot cohort of 33 participants (8 lung cancer, 25 non-cancer controls) underwent oral rinse, bilateral bronchoalveolar lavage (BAL), and bronchial brushing. Microbiome structure was characterized by small subunit ribosomal rRNA gene amplicon sequence variant (ASV) profiling, with indicator-species analysis and SPIEC-EASI correlation-network mapping used to identify ASVs associated with sample type or cancer status.

**Measurements and Main Results:** Sampling method was the dominant variation axis. BAL communities closely resembled oral rinse (64 jointly indicative ASVs; none shared with brushing), whereas bronchial brushing yielded a distinct but low-biomass signal. After excluding host-derived sequences, a single brush-specific indicator ASV affiliated with Sphingomonadaceae was identified within a 17-node co-occurrence module. Cancer-status effects were not detectable, consistent with limited statistical power.

**Conclusions:** Sampling method is the primary determinant of inferred lower respiratory microbiome structure. Bronchial brushing recovers a distinct but low-biomass signal that is obscured when BAL is used in isolation. However, overlap between this low-biomass signal with host and contaminant sequences indicates that confidently resolving a core microbiome will require deeper sequencing and dedicated contamination controls. These methodological findings directly inform the design of future lung cancer- and disease-association studies.

## Introduction

Microbial communities, also known as microbiomes, play wide-ranging roles in health and disease across diverse host systems including human beings [1-4]. The most widely studied environment in the human body is the gastrointestinal tract, where microorganisms contribute to nutrient and drug conversion processes, modulate innate and adaptive immune systems, and influence emotional and cognitive states through the so-called gut-brain axis [5-9]. Dysbiosis in the gut microbiome has been linked to inflammatory bowel disease, type II diabetes, and mood disorders including depression and anxiety [1, 3, 5, 6, 8, 9]. The complex interaction between host and microbiome also extends to cancer, with commensal bacteria directly affecting tumorigenesis, progression and response to therapy [10, 11].

In contrast, although known to be associated with lung disease such as, chronic obstructive pulmonary disease (COPD), cystic fibrosis (CF), and lung cancer, dysbiosis in the lower respiratory tract microbiome remains less well understood [12-14] although evidence suggests that microorganisms inhabiting the lower respiratory tract may be intricately linked to development, progression and treatment response of non-small cell lung cancer [15]. Lung cancer remains the leading cause of cancer-related deaths, with 1.8 million deaths annually and a five-year net survival rate of 22% [16-18]. Critically, lung cancer incidence among individuals traditionally considered low risk, such as people who never smoked (never-smokers), continues to rise in many parts of the world, highlighting the need for improved early preclinical screening methods capable of identifying those who would benefit most from lung cancer screening with low-dose CT scans [19, 20]. This provides both a patient health and point-of-care diagnostic impetus to identify microbial indicators for cancer status.

Low biomass remains a significant barrier to mapping the lower respiratory microbiome and its role in lung health and disease. Bacterial loads in the lower airways are at least two orders of magnitude lower than in the upper respiratory tract [13, 14, 21, 22]. Compounding this sparseness is the presence of significant amounts of human chromosomal and mitochondrial DNA that can confound microbiome sequencing efforts [21, 22]. Despite an increasing focus on lung microbiome research in the context of disease, there is a marked lack of coherence in the accompanying research design. Indeed, contemporary research has involved a variety of sampling and sequencing methods, and different data processing and analysis workflows, complicating efforts to define a coherent atlas of the lower respiratory tract microbiome.

The anatomy of the lung and airway, as well as bidirectional movement of air and moisture, creates ecological gradients, facilitating continuous microbial transport between the external environment, oral cavity, oropharynx, nasopharynx, hypopharynx, and lungs [13, 22, 23]. Previous studies indicate that the oropharynx is the primary source of lung microbiota, with limited evidence for indigenous microorganisms or strong selective pressure within the lungs of healthy individuals, consistent with the adapted island model of lung biogeography [22, 23]. However, these studies were small and performed with proximal brush samples via bronchoscopy, making it technically impossible to avoid oral airway contamination. From a sampling perspective, bronchoscopy-based methods, including bronchoalveolar lavage (BAL) and bronchial brushing, have been used to recover microbial biomass for DNA sequencing [21, 22]. While bronchoalveolar lavage (BAL) is by far the most widely used sampling method in lung microbiome studies [21] such samples have been shown to more closely resemble oral samples than bronchial brushes from the same patient [22], suggesting either carryover from oral secretions as the scope is passed through the oropharynx, or increased signal from greater microbial load within this sample type. Because bronchial brushings represent an observable subset of the respiratory tract microbiome, this form of sampling, targeted distally in ≥4th generation airways, may be more suitable for identifying microbial indicators of dysbiosis and disease progression. and could support more extensive monitoring programs for early lung cancer detection.

Here we chart the community structure of the human lung microbiome, comparing different sampling methods including oral rinse, bilateral bronchoalveolar lavage (BAL) and bilateral bronchial brushings. Multi-level indicator and correlation network analyses were used to test whether bronchial brushing recovers community signal that is distinguishable from BAL and oral rinse, and to assess potential cancer-status effects.

## Methods

Detailed methods related to the participant cohort, sample collection, processing and sequencing are provided in the supplementary text (Supplementary Figures S1 and S2, Supplementary Dataset SD1). Data processing was conducted using ASPIRE: the Amplicon Sequencing Profiler for Investigating Respiratory Ecosystems [24]. This study was approved by the University of British Columbia Research Ethics Board, certificate number H21-00342.

## Results

### Cohort and sampling overview

Within the cohort of 35 enrolled participants, 33 (8 lung cancer patients, 25 non-cancer controls) passed sample-level quality control and were retained for analysis (Table 1). Microbiome profiling was performed on 137 samples (oral rinse (n=32), BAL (n=44), and bronchial brush (n=61)), yielding 7.8, 6.4, and 6.7 million raw paired-end reads, respectively (**Figure 1A**); scope flushes and skin brushes were retained as controls (**Methods**). After applying mitochondrial filtering and a per-sample read threshold, 32 oral rinse, 44 BAL, and 49 bronchial brush samples remained (**Figure 1B**). Bronchial brush samples yielded markedly lower depth (median 1,120 reads versus 19,330 for BAL and 52,546 for oral rinse; per-sample read counts in **Supplementary Dataset SD2**). The three-tier contamination filter (**Methods**) removed 35 of 1,377 ASVs, leaving 1,342 high-confidence ASVs for all downstream comparisons (**Supplementary Dataset SD3)**.

**Table 1.**
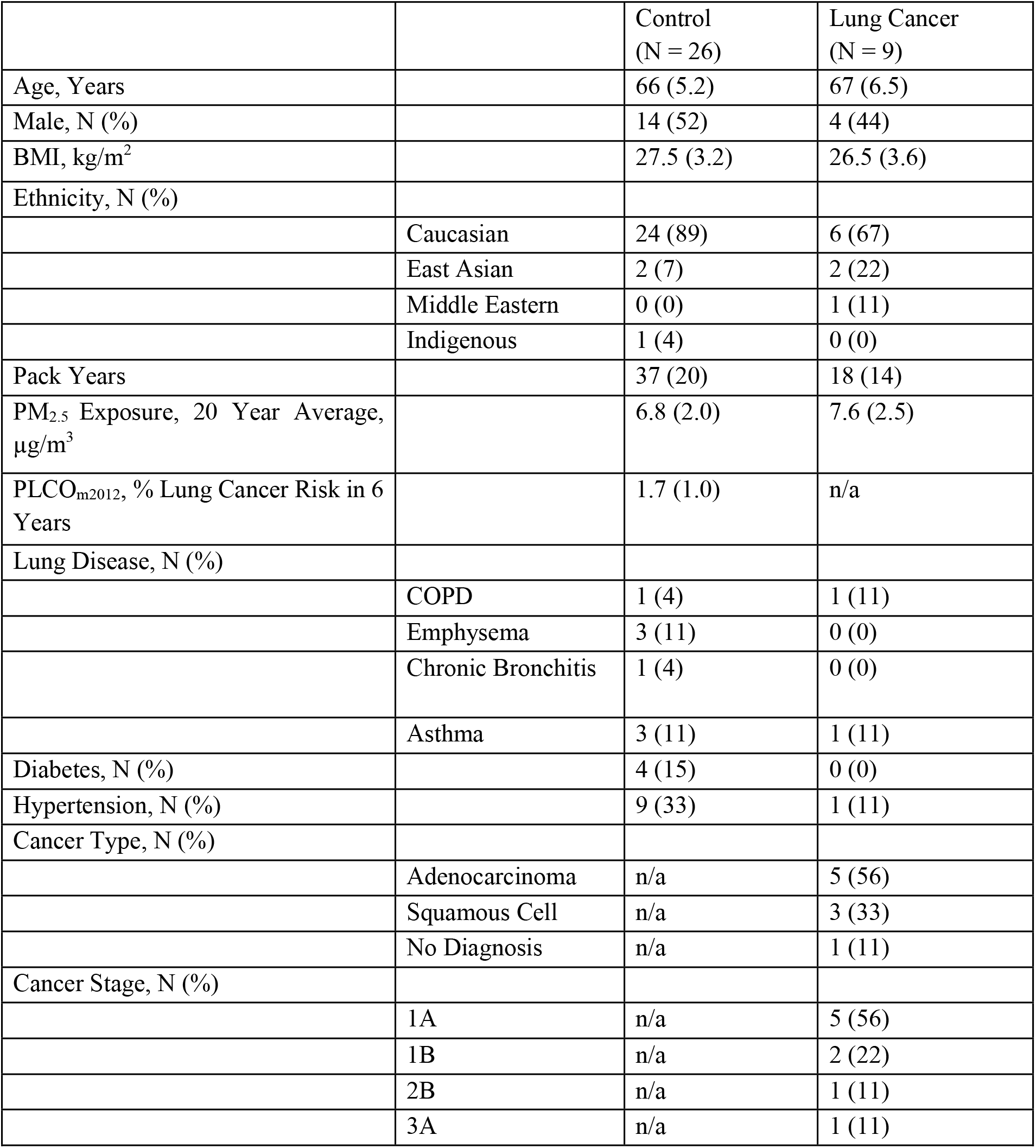
Characteristics of control and lung cancer patients participating in this study. Values are presented as mean (standard deviation) unless specified otherwise. Abbreviations: N (number), Body Mass Index (BMI), One Pack Year is equivalent to smoking one pack of cigarettes per day for a year, Particulate Matter <2.5 µm in diameter (PM_2.5_), Prostate, Lung, Colorectal, Ovarian Cancer lung-cancer risk prediction model 2012 (PLCO_m2012_) [29], Chronic Obstructive Pulmonary Disease (COPD).

**Figure 1.**
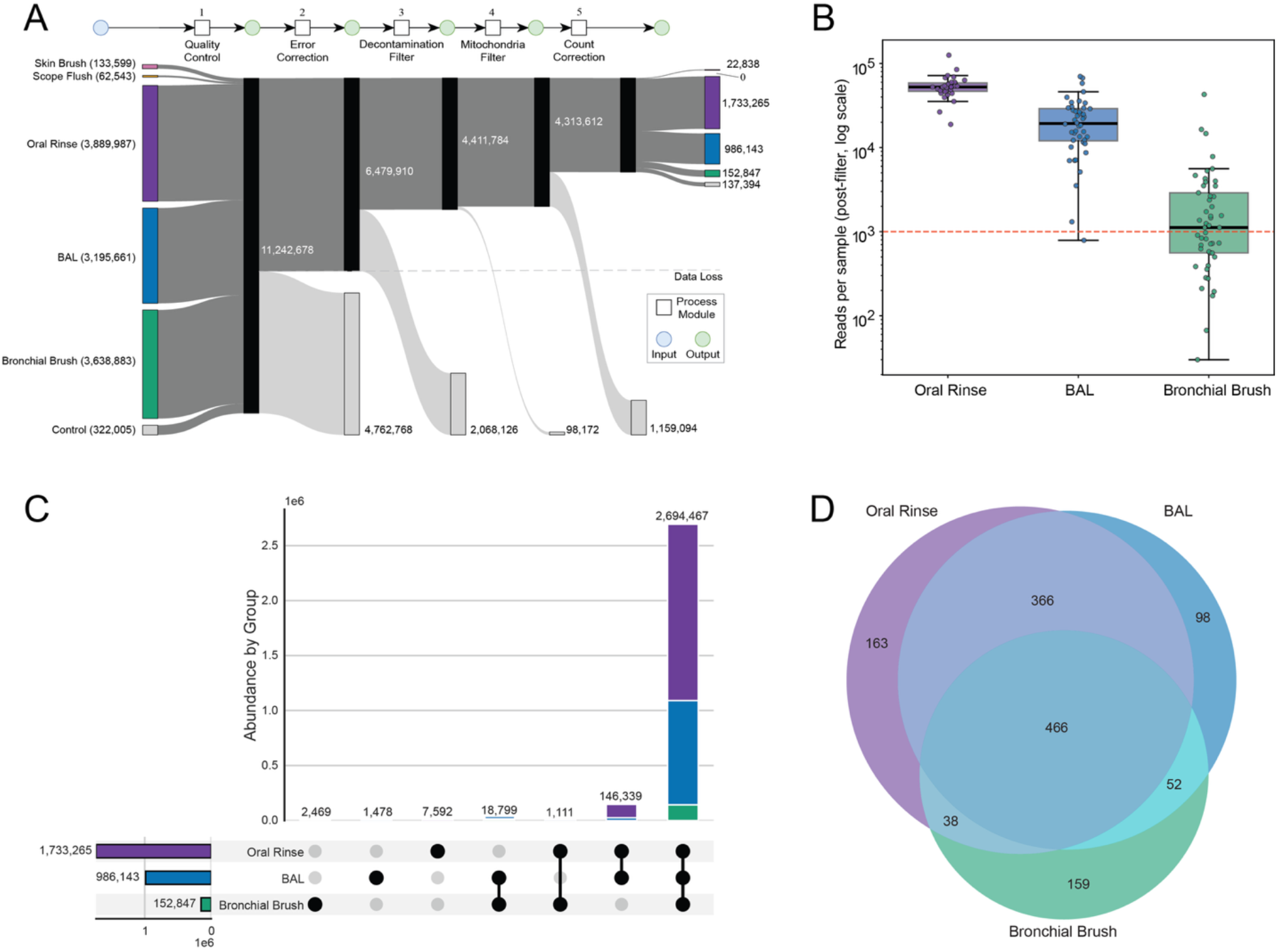
Cohort, sequencing, and ASV sharing across sample types. (A) Total reads retained at each stage of the bioinformatic processing pipeline (raw → quality control → error correction → decontamination → mitochondrial/host filtering), grouped by sample type; the schematic below indicates the corresponding processing modules. All three sample types began with comparable read depth, but bronchial brush samples lost the large majority of reads at the host/mitochondrial-filtering step, reflecting their high host-cell content. (B) Per-sample sequencing depth after filtering (log scale); bronchial brush samples were markedly shallower (median 1,120 reads) than BAL (19,330) or oral rinse (52,546). The dashed line marks the 1,000-read inclusion threshold. (C) UpSet plot of reads associated with ASVs shared among sample types and (D) Venn diagram of ASV sharing: of 1,342 analyzed ASVs, 466 were shared by all three sample types, 366 by oral rinse and BAL only, and 163, 98, and 159 were unique to oral rinse, BAL, and bronchial brush, respectively. The shared-by-all-three ASVs accounted for 2,694,467 of 2,872,255 reads (93.8%), while ASVs unique to a single sample type together accounted for 11,539 reads (0.4%).

In total, 466 ASVs were shared between oral rinse, BAL, and bronchial brush samples, with 366 shared exclusively between oral rinse and BAL samples, 38 ASVs shared exclusively between oral rinse and bronchial brush samples and 52 ASVs shared exclusively between BAL and bronchial brush samples. A total of 163, 98, and 159 unique ASVs were identified in oral rinse, BAL, and bronchial brush samples, respectively (**Figure 1C-D, Supplementary Dataset SD4**). The community structure shared between oral rinse, BAL, and bronchial brush samples was consistent with previous observations that the oral cavity is the primary source of microbes in the lower lung and that other sections of the airway are associated with distinct subsets of microorganisms potentially originating from the oral microbiome [22, 23]. At the same time, the presence of ASVs shared solely between BAL and bronchial brush samples indicates potential for habitat selection.

### Community diversity and taxonomic classification

Alpha and beta diversity metrics provide a high-level overview of microbiome structure. Alpha diversity quantifies the variety of species (as represented by ASVs) within a single sample, encompassing both richness and evenness, and beta diversity measures differences in species composition between samples. Both oral rinse and BAL samples showed significantly higher Shannon alpha diversity than bronchial brush samples, consistent with reduced microbial load within the lower respiratory tract (Wilcoxon signed-rank test; Bronchial Brush vs Oral Rinse: W=13, p=1.3x10^-6^, q=3.9x10^-6^. Bronchial Brush vs BAL: W=45, p=4.7x10^-4^, q=4.7x10^-4^) (**Figure 2A**). Collector’s curves approached saturation for oral rinse and BAL samples, whereas bronchial brush curves continued to rise more steeply and recovered fewer ASVs overall, consistent with incomplete sampling of this low-biomass compartment (**Figure 2B**). Before testing cancer status effects on community structure, we compared cancer-bearing lungs, contralateral lungs (the non-tumor lung in cancer patients), and healthy lungs (**Supplementary Figure S3, Supplementary Table S1, Supplementary Dataset SD5**). Although sample number was limiting, contralateral samples appeared intermediate between cancerous and healthy profiles. To account for this, we excluded contralateral samples from downstream cancer status analyses and compared only cancerous versus healthy lungs. Despite this, Shannon diversity between cancer and non-cancer samples was not significantly different, although more variance between non-cancer participants was observed in bronchial brush samples, indicating increased variation between samples (**Figure 2C)**.

**Figure 2.**
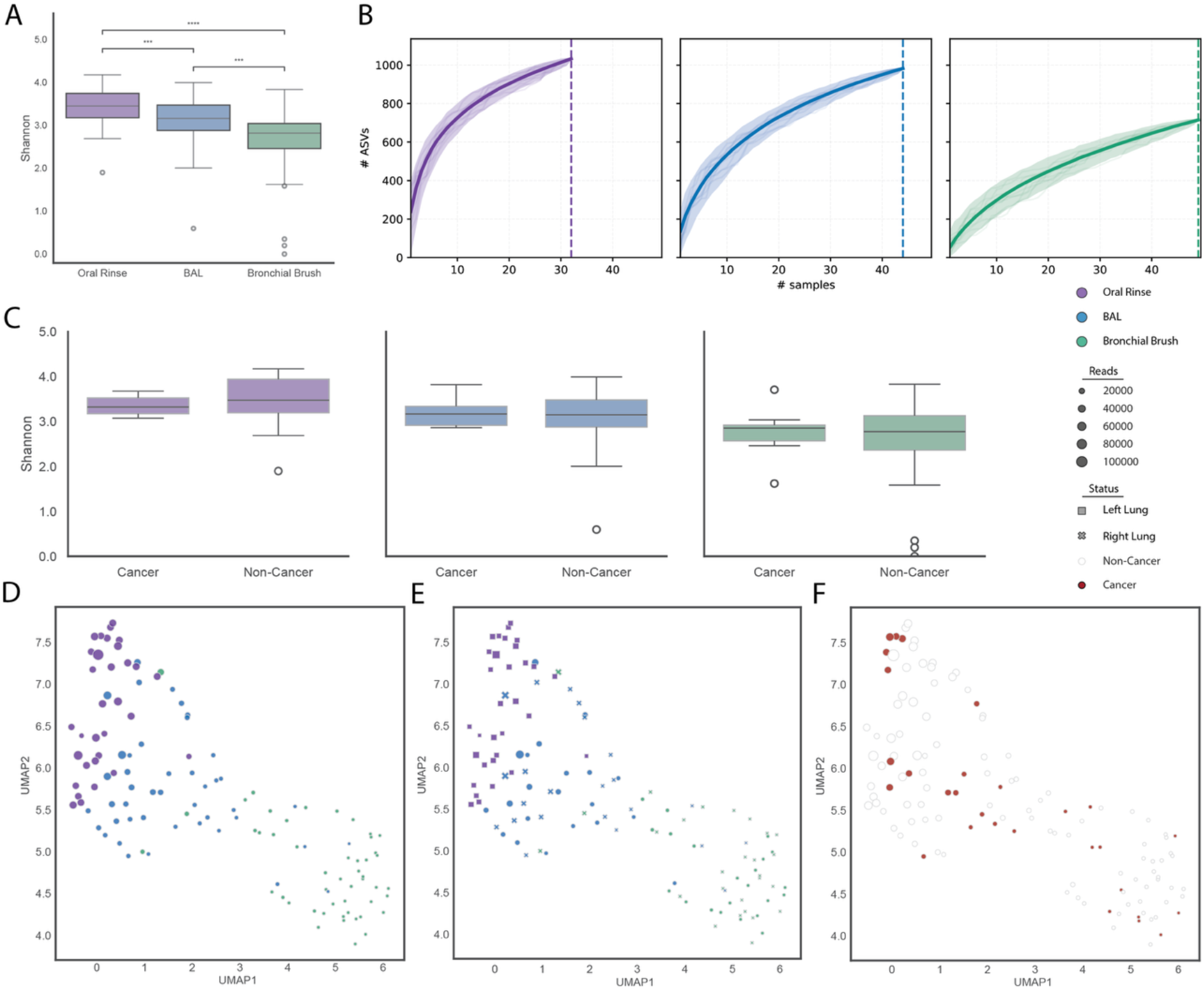
Community diversity by sample type and cancer status. (A) Shannon alpha diversity by sample type; oral rinse and BAL were significantly more diverse than bronchial brush (pairwise Wilcoxon, BH-FDR; ***q<0.001, ****q<0.0001). (B) Collector’s (rarefaction) curves of ASV richness versus number of samples, by sample type; curves approached saturation for oral rinse and BAL but continued to rise for bronchial brush, indicating incomplete sampling of this low-biomass compartment. (C) Shannon diversity by cancer status within each sample type; no comparison was significant. (D–F) UMAP ordination of patient-aware Bray–Curtis dissimilarities (point size scaled by read count): colored by (D) sample type, (E) sample type with lung side indicated, and (F) cancer status. Samples resolved into two broad clusters (low-read-count bronchial brush versus oral rinse and BAL along a read-depth gradient) with no clustering by lung side or cancer status.

Patient-aware pairwise beta diversity analyses showed significant differences between each sample type pair, with the smallest difference observed between bronchial brush and BAL samples and the largest difference between oral rinse and bronchial brush samples. However, a test of multivariate dispersion (PERMDISP) was also significant across sample types (p = 0.001; full diversity statistics in **Supplementary Table S2**), consistent with the greater within-group variance of bronchial brush samples; the sample-type separation detected by PERMANOVA therefore reflects differences in both community composition and dispersion rather than location shifts alone. When visualizing beta diversity using UMAP [25], samples resolved into two broad clusters: one composed predominantly of low-read-count bronchial brush samples, and a second comprising oral rinse and BAL samples arranged along a read-count gradient (**Figure 2D**). Labelling samples within the projection by right or left lung, or by cancer status did not resolve clustering patterns associated with either metadata feature, and beta-diversity distances did not differ significantly by pair type (within-cancer, between-group, or within-non-cancer) (**Figure 2E-F)**. Taken together, these results indicate that microbiome compositions of oral rinse and BAL samples are more closely associated with one another than bronchial brushes, and that bronchial brushes contain a more variable community structure with increased variance relative to oral rinse and BAL samples.

Hierarchical cluster analysis (HCA) resolved two main clusters, one dominated by bronchial brush samples and the other by oral rinse and BAL samples, consistent with the UMAP results (**Figure 3**). Within the oral rinse–BAL cluster, samples partitioned into two subclusters reflecting a preponderance of either oral rinse or BAL samples, while the bronchial brush cluster partitioned primarily by ASV abundance. No specific subclusters were enriched in samples from participants with cancer, suggesting that cancer status alone is unlikely to select for or exclude entire lineages (**Figure 3**). In total, 8 phyla and 38 families differed significantly in relative abundance by sample type (Friedman; q<0.05; host-classified ASVs excluded), including most of the abundant families listed above (Fusobacteriaceae and Lachnospiraceae did not reach significance). Individual pairwise tests between sample types at phylum, family, and genus-level classifications show specific sample type differences (**Figure 4A, Supplementary Dataset SD6 and SD7**). However, no significant differences in taxonomic abundance were observed based on cancer status after multiple-testing correction.

**Figure 3.**
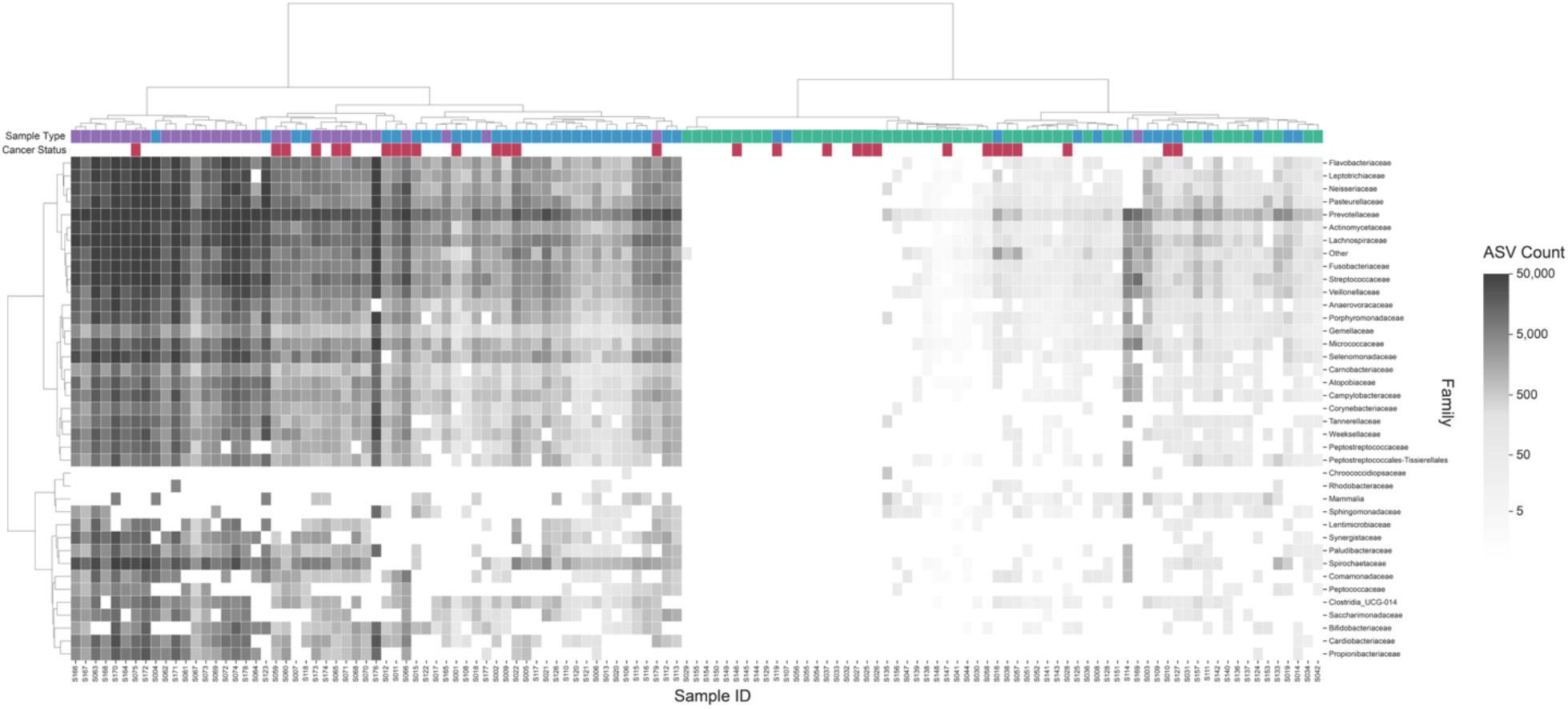
Hierarchical clustering of samples by family-level composition. Hierarchical cluster analysis (Ward linkage, Euclidean distance) of family-level relative-abundance profiles across all analyzed samples. Columns are samples; the annotation bars above the heatmap denote sample type (Oral Rinse, purple; BAL, blue; Bronchial Brush, green) and cancer status (Cancer, red; Non-Cancer, white). Samples resolved into two principal clusters (one dominated by low-read-count bronchial brush samples and the other by oral rinse and BAL samples) with no cluster enriched for cancer status.

**Figure 4.**
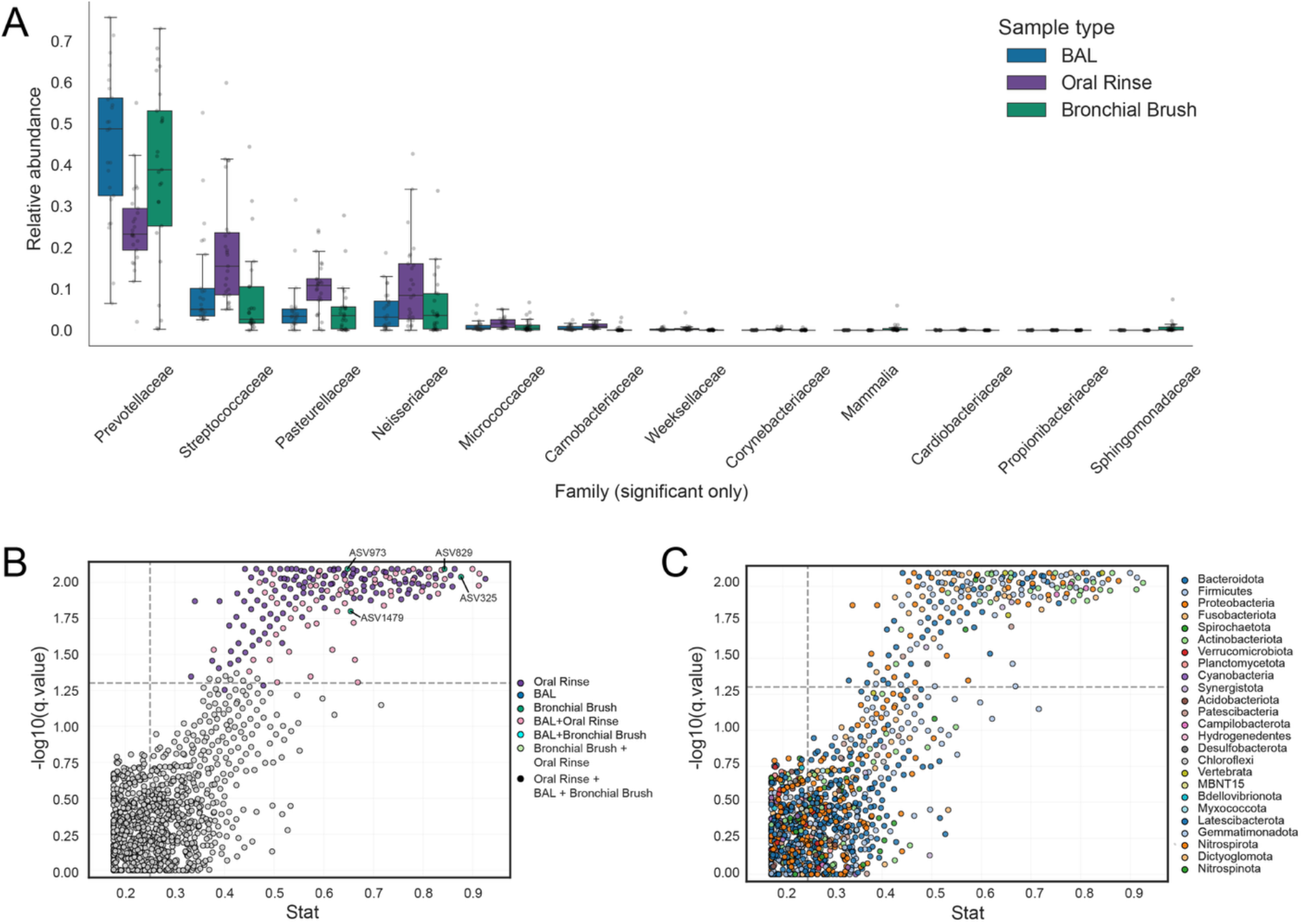
Differential taxonomic abundance and indicator-species analysis. (A) Relative abundance of families that differed significantly among sample types (Friedman, q<0.05), by sample type. (B, C) Multi-level indicator-species analysis: indicator statistic versus ™log10(q-value) for each ASV, colored by (B) the sample-type combination each ASV indicates (dashed lines mark the indicator stat ≥0.25 and q<0.05 thresholds; key bronchial brush–specific ASVs labeled) and (C) phylum. Of 197 significant indicator ASVs, 129 were associated with oral rinse alone, 64 jointly with BAL and oral rinse, and 4 with bronchial brush alone; no ASV jointly indicated BAL and bronchial brush.

Collectively, these results show that most reads mapped to ASVs annotated to 13 families and 13 genera. While these ASVs were primarily shared between oral rinse, BAL, and bronchial brush samples (**Supplementary Results, Supplementary Figure S4, Supplementary Dataset SD4 and SD6)**, their frequency distribution profiles were not uniform, and the abundance of certain taxa varied between sample types. Based on UMAP projection and HCA, BAL samples overlapped with oral rinse samples in ordination space, which suggests bronchial brushes may be more specific in their recovery of core members of the lower respiratory tract microbiome. These results also reinforce the bidirectional nature of the respiratory tract in which microorganisms can migrate or be transported between upper and lower airway intervals while indicating potential for habitat selection.

### Power Analysis

A simulation-based power analysis (**Supplementary Results, Supplementary Figure S5 and Supplementary Table S3**) confirmed that sampling-method comparisons were well powered (>80% at n=10–15), whereas cancer-status comparisons were underpowered at the analyzed cohort size (n=8 cancer participants) across nearly all tests. The cohort size required to reach >80% power depended strongly on the metric: community-level Bray–Curtis comparisons would require approximately 15 cancer participants per sample type, whereas alpha-diversity (Shannon) comparisons remained underpowered even at much larger sizes, reaching 80% only for bronchial brush samples (n≈70) and not at all for BAL or oral rinse up to n=100. This discordance is expected rather than anomalous: alpha diversity collapses an entire community into a single richness/evenness value and is insensitive to compositional shifts that leave overall diversity unchanged, so its low power is consistent with any cancer-associated effect being compositional and subtle rather than a gross change in community diversity. Overall, these power limits frame our interpretation of the null cancer-status findings below.

### Indicator analysis

To investigate potential impacts of niche partitioning and cancer status on lung microbiome structure, a multi-level indicator ASV analysis was performed. This enabled identification of ASVs associated with one or more groups, clusters, or conditions. We first looked for indicator ASVs associated with different combinations of sample types from the set of 1,342 ASVs recovered from oral rinse, BAL, and bronchial brush samples (1,081 ASVs met prevalence thresholds for indicator testing) (**Figure 4C**). In total, 197 ASVs were identified with an indicator score >0.25 and q-value <0.05 (full results in **Supplementary Dataset SD8**) of which 65.5% (n=129) were associated with oral rinse, 32.5% (n=64) with oral rinse + BAL, no ASVs with BAL + bronchial brushes, and 2.0% (n=4) with bronchial brush samples alone (**Supplementary Figure S6**).

The proportion of reads mapping to indicator ASVs varied substantially by sample type association: 2.83% (oral rinse), 11.90% (BAL + oral rinse), 0% (BAL + bronchial brush), and 0.02% (bronchial brush), summing to 14.8% of total dataset reads. Notably, nearly half of the indicator ASVs (91 of 197; 46%) were detected across all three sample types, consistent with differential abundance of widely distributed taxa. ASVs associated with cancer or non-cancer status were also evaluated within each sample type, with no significant indicators surviving multiple-test correction. Statistical power for these comparisons was severely limited at the observed cohort size (status ISA power = 0.001–0.015). After excluding ASVs classified as Vertebrata or Mammalia (host-derived reads that escaped mitochondrial filtering), a single Bronchial Brush–specific indicator ASV was identified, affiliated with Sphingomonadaceae (Sphingobium; stat=0.86, q=0.008). Three further Bronchial Brush–specific indicator ASVs were classified as Vertebrata/Mammalia and excluded as host-derived.

Overall, this analysis identified differentiating features of community structure along the respiratory tract, with many indicators of oral and BAL + oral groups and a select set distinguishing bronchial brushing samples. This confirms the results of prior surveys, which found that lower respiratory tract microbiomes represent subsets of the oral and upper respiratory tract microbiome [22, 23], while also exhibiting distinct, site-specific community structure. The identification of bronchial brush and BAL indicators suggests a distinct and observable community structure present in the lower respiratory tract. This community could be more influenced by disease status, given proximity to cancer sites and greater separation from the external environment.

### Correlation network mapping

Building on the indicator analysis, we next asked whether the habitat-specific signal captured by bronchial brushing is reflected in coordinated co-occurrence patterns among brush-associated ASVs, a more stringent test of community coherence than indicator status alone. A co-occurrence network was constructed using SPIEC-EASI via sparse inverse covariance estimation (glasso), retaining positive partial-correlation edges above an absolute weight threshold. Sample type indicator status, taxonomic affiliation, and module membership were mapped onto the network to identify sampling-method-associated subnetworks.

The significance-thresholded sub-network contained 278 nodes and 703 edges (density = 0.018, average degree = 5.06) distributed across 5 connected components, of which the largest comprised 274 nodes (**Figure 5A–C, Supplementary Dataset SD9**). The network exhibited substantial non-random local clustering (transitivity = 0.237; average local clustering coefficient = 0.194), approximately 11.8-fold elevated over expectation from 1,000 degree-preserving null networks (configuration-model randomization, empirical p < 0.001). Consensus Leiden community detection across multiple resolution parameters (150 replicate runs, consensus threshold 0.95) partitioned the network into modular structure, confirming a robustly modular topology.

**Figure 5.**
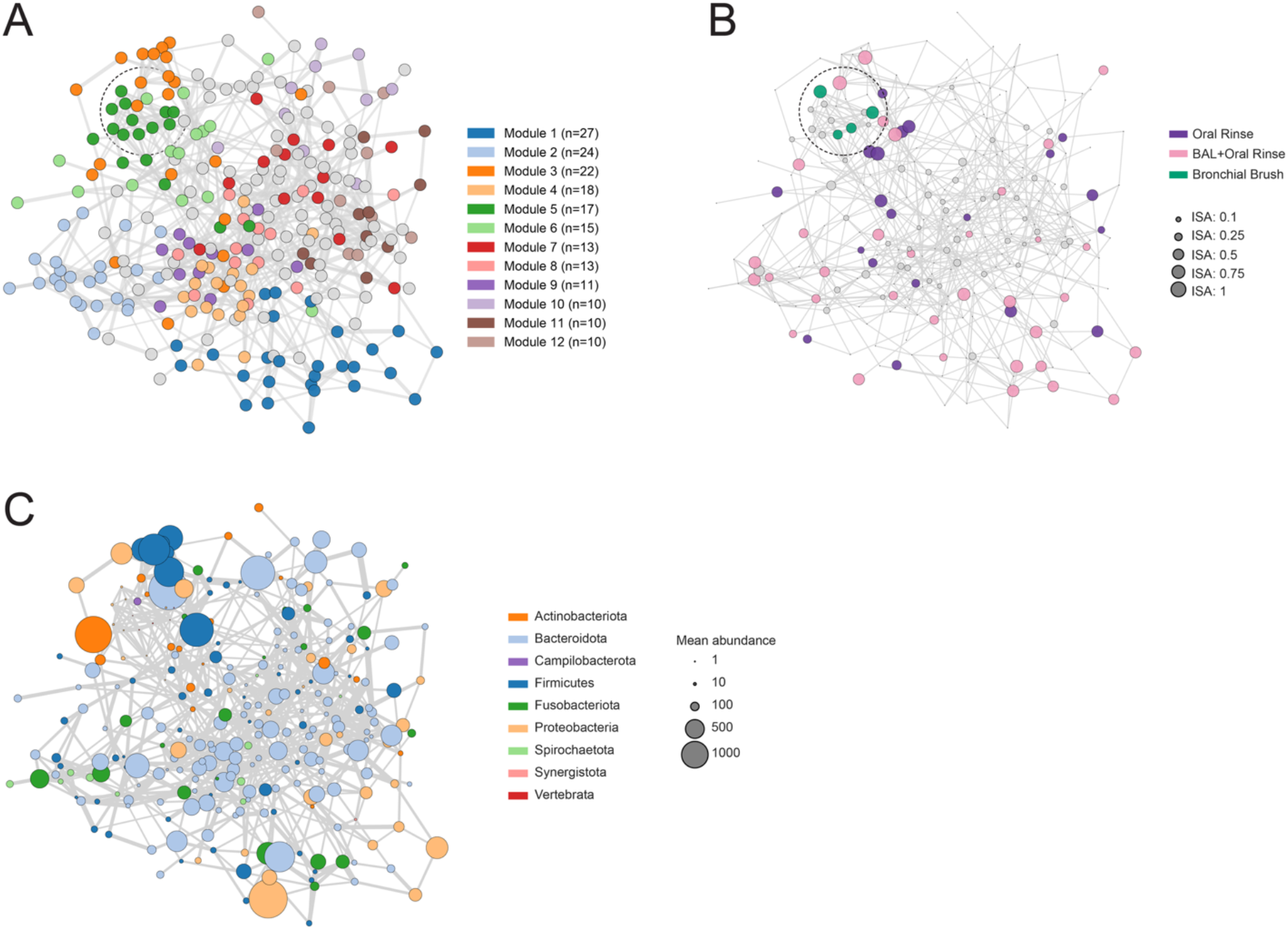
Co-occurrence network structure. SPIEC-EASI co-occurrence network of the analyzed ASVs (278 nodes, 703 edges; edge width scaled by the magnitude of the partial correlation). The same network layout is shown with nodes colored by (A) consensus network module (community), (B) sample-type indicator status (color, indicated sample-type combination; node size, indicator strength), and (C) phylum (node size, mean relative abundance). Both near-significant bronchial brush–specific indicator ASVs co-localized within a single module.

Sample type indicator ASVs distributed non-randomly across modules. A single biologically interpretable bronchial brush–specific indicator ASV (Sphingobium) co-localized within a 17-node network module together with a Steroidobacteraceae ASV that fell just below the significance threshold after correction (p=0.010, q=0.054), and three brush indicators identified as host contamination (**Figure 5, Supplementary Dataset SD9**). Beyond these indicators, the module comprised 11 additional bacterial ASVs (predominantly Proteobacteria) and an additional host-classified (Homo sapiens) ASV (full membership and taxonomy in **Supplementary Dataset SD9**; cross-referenced against published reagent and laboratory contaminant lists in **Supplementary Table S4**). The co-localization of the brush indicators with a near-significant Steroidobacteraceae ASV within a single module supports the interpretation that bronchial brushing recovers a community ecologically distinct from that captured by BAL or oral rinse, despite both originating from bronchoscopy-based sampling of the lower airway.

In contrast, the 64 BAL + oral rinse co-indicator ASVs distributed less coherently. Only 35 of 64 (55%) met the prevalence threshold for inclusion in the co-occurrence network; among those, the largest single concentration was 9 indicators within one 30-node module, with the remainder dispersed across 11 modules. The 129 oral rinse–only indicators were even more diffusely organized: only 20 (16%) appeared in the network at all (consistent with their lower per-ASV prevalence across the cohort), and these were distributed across 7 modules, with no more than 7 indicators in any single module. The diffusive tendency of upper-airway-derived indicators contrasts sharply with the focal concentration of both bronchial brush–associated indicators within a single module, reinforcing the interpretation that bronchial brushing recovers an ecologically distinct microbial signal that is both detectable as habitat-specific indicators and reproducible as a coherent co-occurrence module.

## Discussion

In this study, sampling method emerged as the dominant tested axis of variation in respiratory microbiome structure, substantially exceeding cancer-status effects. Across three convergent lines of evidence (patient-aware Bray-Curtis PERMANOVA, multi-level indicator-species analysis, and SPIEC-EASI co-occurrence network mapping), sample type consistently differentiated communities recovered by oral rinse, BAL, and bronchial brushing. These analyses are not fully independent, as all derive from the same compositional ASV table and are driven by the same dominant sample-type structure, and the network is constructed from the pooled cohort such that co-occurrence of brush-associated ASVs partly reflects their shared presence in brush samples.

BAL samples, despite originating from bronchoscopy-based sampling distal to the oropharynx, clustered closely with oral rinse samples in both UMAP projection and hierarchical clustering and shared 64 jointly indicative ASVs with oral rinse compared to zero co-indicators with bronchial brushing. This convergence indicates that BAL captures a mixed signal reflecting the lower airway with substantial oral carryover during scope passage through the oropharynx. This is consistent with prior reports describing BAL composition as more closely resembling oral than bronchial-brush communities from the same patient [22, 23]. The dominance of oral cavity-affiliated families including Prevotellaceae, Streptococcaceae, Veillonellaceae, and Fusobacteriaceae among the BAL+oral rinse co-indicators reinforces this interpretation. From a study-design perspective, BAL provides a higher-biomass and more readily recoverable sample, but the analyses presented here indicate that its compositional similarity to oral microbiota may attenuate signals originating specifically in the lower airway.

In contrast, bronchial brushing yielded a small but focally coherent signal. Two habitat-specific indicator ASVs, affiliated with Sphingomonadaceae and Steroidobacteraceae, co-localized within a single co-occurrence module alongside additional ASVs characteristic of low-biomass airway sampling. Two complementary lines of evidence, including indicator analysis identifying these ASVs as habitat-preferential, and SPIEC-EASI community detection assigning them to the same module, resolved the same taxa, pointing to a coherent non-random co-occurrence pattern. Notably, ASVs affiliated with Sphingomonadaceae have been reported as low-abundance residents in airway samples from healthy and diseased individuals [26, 27]. The separation of both brush indicator ASVs into a module distinct from the BAL- and oral-rinse-associated community suggests that bronchial brushing recovers community signal at least partly distinct from oropharyngeal carryover, and that this signal is sufficiently structured to be detectable by network-based methods even at the modest sample sizes characteristic of bronchoscopy-based pilot studies.

Bronchial brush samples in this cohort yielded substantially lower per-sample sequencing depth than other sample types (median 1,120 reads versus 52,546 for oral rinse and 19,330 for BAL) and showed a higher rate of phylum-unclassified reads (∼23%), both factors that amplify the relative contribution of any low-level contamination signal. Four negative extraction-control samples were sequenced and incorporated into the contamination-filtering steps described in Methods (`decontam` prevalence and within-sample-type frequency methods), which, together with a biological-plausibility screen for thermophilic environmental taxa, removed 35 ASVs. Definitive separation of residual lower-airway signal from procedural background will nonetheless benefit from deeper negative-control sequencing combined with increased sampling depth and host-DNA depletion to reduce mitochondrial background.

Although cancer-status effects were not discerned, the methodological findings of this study have direct implications for future research. Power analysis indicated that status indicator-species comparisons were severely underpowered at the analyzed cohort size (status ISA power 0.001-0.015); Future studies investigating cancer-associated microbial signatures will require cohorts of at least 15 participants with lung cancer per sample type to achieve adequate power to detect community-level differences, with even larger cohorts needed for sample-type-stratified analyses at the family-abundance level. An additional challenge is the recruitment of sufficiently large cohorts of healthy individuals for comparison with patients with lung cancer, particularly given the invasive nature of lower airway sampling. The null findings with respect to cancer status are consistent with a recent large study of 940 never-smokers with lung cancer, which similarly reported no clinically relevant microbial associations [35]. Importantly, however, that study did not include healthy non-cancer controls, limiting its ability to evaluate microbiome differences between healthy individuals and patients with lung cancer.

In conclusion, sampling method was observed to play a primary role in determining lung microbiome structure. Future brush studies integrating expanded sampling with host DNA depletion are needed to provide both the necessary coverage for genome-resolved analyses [28] and increased statistical power for identifying cancer-associated microbial signals supporting biomarker discovery.

## Supporting information

SPARK_supplementary_material

## Author Contributions

Renelle Myers, Crista Bartolomeu, Scott A. Borden, Stephen Lam developed the cohort study, conducted bronchoscopy experiments, and prepared participant samples and metadata. Avery J. C. Noonan extracted DNA from participant samples and coordinated library production and sequencing. Ryan McLaughlin, Anika Nag, Steven Chen and Steven J. Hallam contributed to data processing and visual analysis workflow development. Avery J. C. Noonan, Ryan McLaughlin, Anika Nag, and Steven Chen processed sequence data. Avery J. C. Noonan conducted the power analysis with input from Renelle Myers, Crista Bartolomeu, Scott A. Borden, Stephen Lam. Avery J. C. Noonan, Ryan McLaughlin, Anika Nag, Steven Chen, and Steven J. Hallam conducted primary data analysis and interpretation. Renelle Myers, Crista Bartolomeu, Scott A. Borden, Stephen Lam provided clinical perspectives on sample collection and data interpretation. Avery J. C. Noonan, Ryan McLaughlin, Anika Na, Steven Chen and Steven J. Hallam prepared the first draft of the manuscript. All authors contributed to the revision of the draft manuscript and approved the final draft of the manuscript.

## Supplementary Material

Supplementary Results, Figures and Tables are provided in the associated Supplementary Material. Supplementary Datasets (SD1–SD10) are available via Zenodo (DOI: 10.5281/zenodo.22839013).

## Acknowledgements

We would like to thank Tom Pfeifer for help in establishing automation workflows in the Biofactorial automation core facility.

## Data Availability

The filtered ASV count table and all large per-feature data files are available as Supplementary Datasets (SD1–SD10) via Zenodo (DOI: 10.5281/zenodo.22839013).

## References

1. Hou, K., et al., Microbiota in health and diseases. Signal Transduct Target Ther, 2022. 7(1): p. 135.

2. Aggarwal, N., et al., Microbiome and Human Health: Current Understanding, Engineering, and Enabling Technologies. Chem Rev, 2023. 123(1): p. 31–72.

3. VanEvery, H., et al., Microbiome epidemiology and association studies in human health. Nature Reviews Genetics, 2023. 24(2): p. 109–124.

4. Human Microbiome Project, C., Structure, function and diversity of the healthy human microbiome. Nature, 2012. 486(7402): p. 207–14.

5. Carabotti, M., et al., The gut-brain axis: interactions between enteric microbiota, central and enteric nervous systems. Ann Gastroenterol, 2015. 28(2): p. 203–209.

6. Sorboni, S.G., et al., A Comprehensive Review on the Role of the Gut Microbiome in Human Neurological Disorders. Clinical Microbiology Reviews, 2022. 35(1).

7. Crouwel, F., H.J.C. Buiter, and N.K. de Boer, Gut Microbiota-driven Drug Metabolism in Inflammatory Bowel Disease. Journal of Crohn’s and Colitis, 2021. 15(2): p. 307–315.

8. Fan, Y. and O. Pedersen, Gut microbiota in human metabolic health and disease. Nature Reviews Microbiology, 2021. 19(1): p. 55–71.

9. Valdes, A.M., et al., Role of the gut microbiota in nutrition and health. BMJ, 2018. 361: p. k2179.

10. C, d.M., et al., Global burden of cancers attributable to infections in 2008: a review and synthetic analysis - PubMed. The Lancet. Oncology, 2012 Jun. 13(6).

11. E, E., et al., Inflammation-induced cancer: crosstalk between tumours, immune cells and microorganisms - PubMed. Nature reviews. Cancer, 2013 Nov. 13(11).

12. Natalini, J.G., S. Singh, and L.N. Segal, The dynamic lung microbiome in health and disease. Nature Reviews Microbiology, 2023. 21(4): p. 222–235.

13. Dickson, R.P. and G.B. Huffnagle, The Lung Microbiome: New Principles for Respiratory Bacteriology in Health and Disease. PLoS Pathog, 2015. 11(7): p. e1004923.

14. Li, R.M., J. Li, and X.K. Zhou, Lung microbiome: new insights into the pathogenesis of respiratory diseases. Signal Transduction and Targeted Therapy, 2024. 9(1).

15. F, W., et al., The Lung Microbiome: A Central Mediator of Host Inflammation and Metabolism in Lung Cancer Patients? - PubMed. Cancers, 12/22/2020. 13(1).

16. Cancer, I.A.f.R.o. Global Cancer Observatory: Cancer Today. 2022; Available from: https://gco.iarc.fr/today.

17. Canada, C.C.S.A.C.C.C.S.S.C.P.H.A.o., Canadian Cancer Statistics 2025. 2025.

18. Jeon, J., et al., Smoking and Lung Cancer Mortality in the United States From 2015 to 2065: A Comparative Modeling Approach. Ann Intern Med, 2018. 169(10): p. 684–693.

19. Tseng, C.H., et al., The Relationship Between Air Pollution and Lung Cancer in Nonsmokers in Taiwan. J Thorac Oncol, 2019. 14(5): p. 784–792.

20. Sun, S., J.H. Schiller, and A.F. Gazdar, Lung cancer in never smokers--a different disease. Nat Rev Cancer, 2007. 7(10): p. 778–90.

21. Carney, S.M., et al., Methods in Lung Microbiome Research. American Journal of Respiratory Cell and Molecular Biology, 2020. 62(3): p. 283–299.

22. Dickson, R.P., et al., Bacterial Topography of the Healthy Human Lower Respiratory Tract. Mbio, 2017. 8(1).

23. Dickson, R.P., et al., Spatial Variation in the Healthy Human Lung Microbiome and the Adapted Island Model of Lung Biogeography. Annals of the American Thoracic Society, 2015. 12(6): p. 821–830.

24. McLaughlin, R.J., et al., ASPIRE: the Amplicon Sequencing Profiler for Investigating Respiratory Ecosystems. bioRxiv, 2026.08.12.743000; doi :10.64898/2026.08.05.743000

25. McInnes, L., et al., UMAP: Uniform Manifold Approximation and Projection. Journal of Open Source Software, 2018/09/02. 3(29).

26. Zakharkina, T., et al., Analysis of the Airway Microbiota of Healthy Individuals and Patients with Chronic Obstructive Pulmonary Disease by T-RFLP and Clone Sequencing. PLOS ONE, Jul 9, 2013. 8(7).

27. Jin, C., et al., Commensal Microbiota Promote Lung Cancer Development via γδ T Cells. Cell, 2019/02/21. 176(5).

28. Kieft, B., et al., Genome-resolved correlation mapping links microbial community structure to metabolic interactions driving methane production from wastewater. Nat Commun, 2023. 14(1): p. 5380.

29. Tammemägi Martin, C., et al., Selection Criteria for Lung-Cancer Screening. New England Journal of Medicine, 2013. 368(8): p. 728–736.

