## Supplementary material for "Methodological Impacts on Microbiome Structure and Indicator Status in the Human Lower Respiratory Tract": SPARK_supplementary_material

### 1. Supplementary Methods

#### *1.1 Participant Cohort*

A prospective cohort of 35 participants were enrolled: 26 non-cancer controls and 9 participants with newly diagnosed early-stage (1A–3A) lung cancer. Non-cancer controls were former-tobacco users considered high-risk for lung cancer via the PLCOm2012 model (six-year risk >1.5%) [26] and/or US Preventive Services Task Force 2013 criteria (>30 pack-years, age 50–80), enrolled through the British Columbia lung cancer screening trial. Each had undergone a low-dose chest CT within the prior year showing no evidence of lung cancer or pulmonary abnormality. Cancer participants were former-tobacco users (defined as ≥1 year of cessation), recruited through the thoracic surgery clinic at Vancouver General Hospital and the interventional respiratory clinic at BC Cancer. Exclusion criteria were antibiotic use within the prior three months or current inhaled corticosteroid use. Following sample-level quality-control filtering (see Methods), 2 participants (1 cancer, 1 control) were excluded, yielding a final analyzed cohort of 33 participants (8 cancer, 25 non-cancer).

#### *1.2 Sample Collection*

A total of 200 samples, encompassing scope rinses, oral rinse, BAL, bronchial brushes, and skin brushes, were collected from study cohort participants. Participants with lung cancer undergoing diagnostic or staging bronchoscopy first performed an oral rinse with 10 ml of sterile saline, performing a gargle-swish-gargle for 10 seconds, then expectorated into a sterile sample container. A skin brushing was then performed with a cytology brush applied to the antecubital fossa with 10 strokes, and placed in cytolyt. Prior to the bronchoscopy, the working channel of the bronchoscope was flushed with 10 ml of sterile saline and collected into a sterile container from the distal tip of the scope working channel. During the bronchoscopy, protected specimen bronchial brushings were taken as close to the tumor as possible, guided by a radial ultrasound probe to identify the tumour, and passed through the radial guide sheath within the working channel of the bronchoscope. Samples from the corresponding segment within the contralateral non-tumour lung were completed, followed by a BAL with the distal tip of the scope wedged into the subsegment leading to the lesion. 60 ml of sterile saline was instilled, and 20 ml of return fluid was collected in a sterile sample container. A BAL was then repeated in the matching segment in the contralateral lung. Control participants undergoing bronchoscopy for the study also performed an oral rinse and the bronchoscope was flushed with sterile saline and collected prior to procedure as described above. Bronchial brushings were performed through the working channel of the bronchoscope in the right (RB1) and left upper lobe (LB1+2) most apical segments. A BAL was then performed as described above, in these segments, bilaterally if the participant was clinically tolerating the procedure. For each participant, brushing and skin swab samples were submerged in 1ml Cytolyt fixative, transported on dry ice, and ultimately stored at -80 °C until DNA extraction.

#### *1.3 DNA Extraction and SSU rRNA Gene Amplicon Sequencing*

Microbiome structure was determined using small subunit ribosomal (SSU rRNA) gene amplicon sequencing. DNA was extracted from the sample types described above using the ZymoBIOMICS MagBead DNA/RNA kit, following manufacturers specifications, in the Biofactorial automation core facility at UBC (British Columbia, CA). SSU rRNA amplicon sequencing was performed by the Biofactorial high-throughput biology facility (University of British Columbia). Sequencing was performed using Earth Microbiome Project (EMP) primers 515F (5' GTGYCAGCMGCCGCGGTAA 3') and 926R (5' CCGYCAATTYMTTTRAGTTT 3') which capture the V4 and V5 variable regions of the SSU rRNA gene. Sequencing was performed on an Illumina NovaSeq 6000 platform, using the 250-bp paired-end kit (V2 500-cycle PE Chemistry, Illumina Inc, San Diego, CA, USA).

#### *1.4 SSU rRNA Gene Amplicon Sequence Processing*

Sample data was processed using a bespoke sequence analysis workflow implementing a series of computational steps organized into 5 modules related to 1) quality control, 2) error correction, 3) decontamination, 4) mitochondrial sequence filtering, and 5) count correction using negative and non-target controls, followed by data analysis and visualization (**Supplementary Figure S1**). Samples with fewer than 1,000 read counts after mitochondrial filtering and decontamination were excluded; participants with no remaining samples after this filter were excluded from analysis. This threshold was applied to raw counts at the working-table stage. Sequences without a taxonomic assignment were then removed, and a count-correction step adjusted per-ASV counts against negative and non-target controls. Because both steps act after the read threshold, per-sample totals in the analyzed dataset can fall below 1,000: 24 of 125 samples (23 bronchial brush, one BAL) carry fewer than 1,000 corrected reads, and 10 carry fewer than 500 (Supplementary Dataset SD2). These corrected counts are the high-confidence read set used for all downstream analyses.

*1.5 Contamination Filtering*

Following the core processing workflow, a three-tier contamination filter was applied to the analyzed ASV set. First, the `decontam` prevalence method (Davis et al. 2018) was applied to the pooled sample set against four negative extraction controls (PBS, PBS_twz, Negative_96, and Negative_man), flagging ASVs significantly enriched in negative controls relative to biological samples ($p < 0.1$). Second, the `decontam` frequency method was applied separately within each sample type, flagging ASVs whose relative abundance was inversely correlated with DNA concentration within at least one sample type ($p < 0.1$); stratifying by sample type avoids confounding the structurally lower DNA yield of bronchial brush samples with genuine contamination. Third, a biological-plausibility screen removed ASVs assigned to taxa with growth optima incompatible with human airway physiology (thermophilic and environmental lineages including Coprothermobacterota, Thermotogota, Caldatribacteriota, and methanogenic Euryarchaeota), which are not detectable by either `decontam` method because they are absent from extraction blanks and arrive via run-level index cross-talk rather than reagent contamination. Across the three tiers, 35 of 1,377 ASVs were removed, yielding the final set of 1,342 ASVs. Removed ASVs and the filter responsible for each are listed in **Supplementary Dataset SD1;** `decontam` score distributions and negative-control comparisons are shown in **Supplementary Figure S2**.

*1.6 Statistical Analysis*

Alpha diversity (Shannon index) was compared between sample types using patient-aware paired Wilcoxon tests. Beta diversity (Bray–Curtis) was assessed by patient-blocked PERMANOVA (999 permutations, seed 42), with PERMDISP evaluated alongside to test dispersion homogeneity. Differential taxon abundance across sample types was tested by Friedman omnibus test with paired Wilcoxon post-hoc comparisons; because the Friedman test requires complete blocks, these comparisons were restricted to the 25 participants who contributed paired oral rinse, BAL, and bronchial brush samples. Indicator ASVs were identified using multipatt (R package indicspecies)

[27], retaining ASVs with indicator stat $\geq 0.25$ and $q < 0.05$. Co-occurrence networks were constructed with SPIEC-EASI [28] on ASVs present in the network-input set, with model selection by StARS [29]. Statistical power was estimated by simulation: for each comparison, additional samples were generated across a range of cohort sizes under the effect sizes observed in this cohort, and power was taken as the proportion of simulated datasets yielding a significant result. The Benjamini–Hochberg false-discovery-rate correction was applied throughout.

### 2. Supplementary Results

#### *2.1 Taxonomic relative abundance*

We next compared taxonomic profiles at the phylum and family level, exploring whether abundance of these taxonomic groups differed significantly by sample type or cancer status. Both were dominated by taxa typical of the oral and upper respiratory tract: the most abundant phyla were Bacteroidota, Firmicutes, Proteobacteria, Fusobacteriota, and Actinobacteriota, and the most abundant families included Prevotellaceae, Streptococcaceae, Neisseriaceae, Pasteurellaceae, and Veillonellaceae (full relative abundances in **Supplementary Dataset SD6**). At the phylum level, reads mapping to ASVs associated with Bacteroidota, Firmicutes, Proteobacteria, Fusobacteriota, and Actinobacteriota were the most abundant across the sample set (Supplementary Figure S4A;Supplementary Dataset SD6), representing on average 41.4%, 27.6%, 17.3%, 9.0%, and 3.7%, respectively. In total, 8 phyla showed significant differences in relative abundance between sample types (Friedman; q<0.05; Vertebrata-classified ASVs excluded as host-derived;Supplementary Dataset SD7a), including four of the five most abundant phyla (Bacteroidota, Firmicutes, Proteobacteria, Actinobacteriota; Fusobacteriota did not reach significance, q=0.20), as well as less-abundant Phyla including Campilobacterota, Patescibacteria, and Synergistota. At the family level, reads mapping to ASVs associated with Prevotellaceae, Streptococcaceae, Neisseriaceae, Pasteurellaceae, Veillonellaceae, Fusobacteriaceae, Porphyromonadaceae, Leptotrichiaceae, Actinomycetaceae, Flavobacteriaceae, Lachnospiraceae, Micrococcaceae, and Gemellaceae were the most abundant across the sample set (Figure 3;Supplementary Dataset SD6), representing on average 33.5%, 15.0%, 8.9%, 8.1%, 7.7%, 7.4%, 5.4%, 1.6%, 1.4%, 1.3%, 1.2%, 1.2%, and 1.1%, respectively or 93.7% of total reads mapped. In multiple instances, the frequency distribution of reads mapping to ASVs varied between sample types.

*2.2 Power Analysis*

Given the pilot-scale cohort used in the study, we conducted a systematic power analysis to evaluate the effect sizes and sampling depths required to identify significant differences in ASV abundance and community structure between sample types (the primary analytical focus of this study) and between cancer-status groups (a secondary comparison constrained by pilot-scale cohort size). Effect-size estimates derived from underpowered cohorts are subject to inflation; observed power should therefore be interpreted as an upper bound, and projected sample requirements as a lower bound for future studies. Power analysis of sample-type comparisons indicated that our study design was highly sensitive to differences between sampling methods. For beta diversity (Bray–Curtis PERMANOVA), statistical power exceeded 80% for all sample-type comparisons at n=15 (omnibus power approaching 100% at n=10;Supplementary Figure S5a). Alpha diversity (Shannon index) power crossed the 80% threshold at approximately n=15 (0.96); at n=10, Shannon power was 0.78, indicating that beta diversity is the more sensitive marker of sampling-method effects in this design. The strong signal across both metrics reflects the ecological partitioning of microbiomes recovered from oral rinse, BAL, and bronchial brushing, and indicates that the sampling-method effects reported in this study are robust and likely reflective of true biological signal.

Statistical sensitivity for sample-type comparisons varied by taxonomic resolution (Supplementary Figure S5b). Phylum-level analyses exhibited limited power at small cohort sizes, requiring approximately 30 cancer participants in BAL samples, with Oral Rinse phylum-level comparisons reaching 80% power at a similar cohort size. Family-level comparisons were more sensitive, particularly in BAL samples where 10 cancer participants were sufficient to reach the 80% threshold. Family-level comparisons in bronchial brush samples required larger cohorts (n=15 cancer participants for 80% power), reflecting smaller per-family effect sizes within this sampling method. In contrast, comparisons between cancer and non-cancer participants were underpowered across nearly all statistical tests at the analyzed cohort size (n=8 cancer participants), with the sole exception of community-level Bray–Curtis PERMANOVA in bronchial brush samples (power = 0.85). Beta-diversity simulations parameterized by the observed effect sizes indicated that approximately 15 cancer participants per sample type would achieve >80% power for community-level Bray–Curtis comparisons (BAL n=15, oral rinse n=15, bronchial brush n=8). Family-level abundance comparisons required modestly larger cohorts (n=10–20 depending on sample type) and phylum-level comparisons spanning a wider range (n=10 for bronchial brush to n=30 for oral rinse). In marked contrast, alpha-diversity (Shannon) comparisons remained underpowered across a far wider range: power reached 80% only for bronchial brush samples (at n≈70) and did not reach 80% for BAL (maximum 0.07) or oral rinse (maximum 0.76) even at n=100. Because alpha diversity summarizes a whole community as a single richness/evenness value, it is insensitive to compositional shifts that do not change overall diversity; its low power is therefore consistent with any cancer-associated effect being compositional and subtle rather than a gross diversity change and should not be read as evidence against a detectable cancer signal in adequately powered designs. These power limitations inform our interpretation of the null cancer-status findings.

### 3. Supplementary Figures

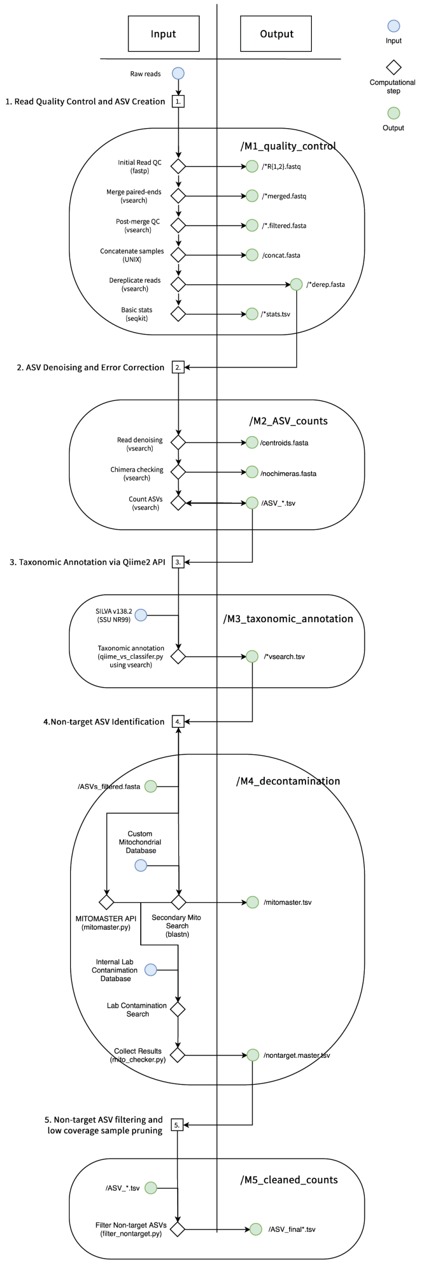


**Supplementary Figure S1. Bioinformatic processing workflow. Schematic** of the sequence-analysis pipeline used to generate the analyzed ASV dataset, organized into five modules: (M1) read quality control and ASV creation (paired-end QC, merging, and dereplication with fastp and VSEARCH); (M2) ASV denoising and error correction (UNOISE denoising, UCHIME chimera removal, and ASV quantification); (M3) taxonomic annotation against the SILVA reference database (QIIME 2 / VSEARCH); (M4) decontamination and non-target sequence identification, including mitochondrial and host-sequence detection (MITOMASTER, BLAST) and laboratory-contaminant screening; and (M5) non-target ASV filtering and removal of low-coverage samples, yielding the final cleaned count table. For each step, the computational tool is indicated in parentheses and the corresponding output file is shown on the right. Input nodes are shown in blue, computational steps as diamonds, and output files in green.

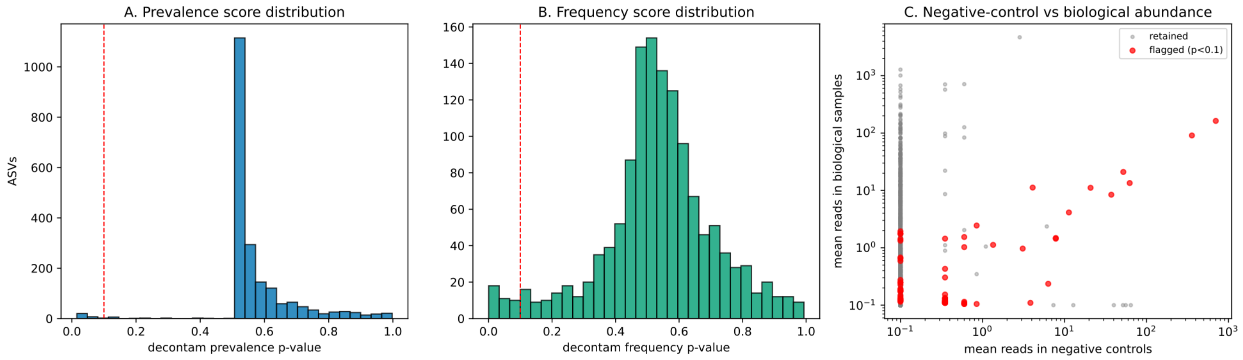


**Supplementary Figure S2. Contamination-filter diagnostics.** Diagnostic distributions from the decontam-based contamination screen applied to the full ASV set (n = 3,174 ASVs scored against four negative extraction controls). (A) Distribution of decontam prevalence-method p-values; (B) distribution of decontam frequency-method p-values; the dashed line marks the $p < 0.1$ flagging threshold used in both methods. (C) Mean per-ASV read abundance in negative-control versus biological samples (log scale); ASVs flagged as putative contaminants ($p < 0.1$) are highlighted in red. Across the three-tier filter (prevalence, within-sample-type frequency, and biological-plausibility screen), 65 ASVs were flagged in total, of which 37 were present in the analyzed table and removed (Supplementary Dataset SD1).

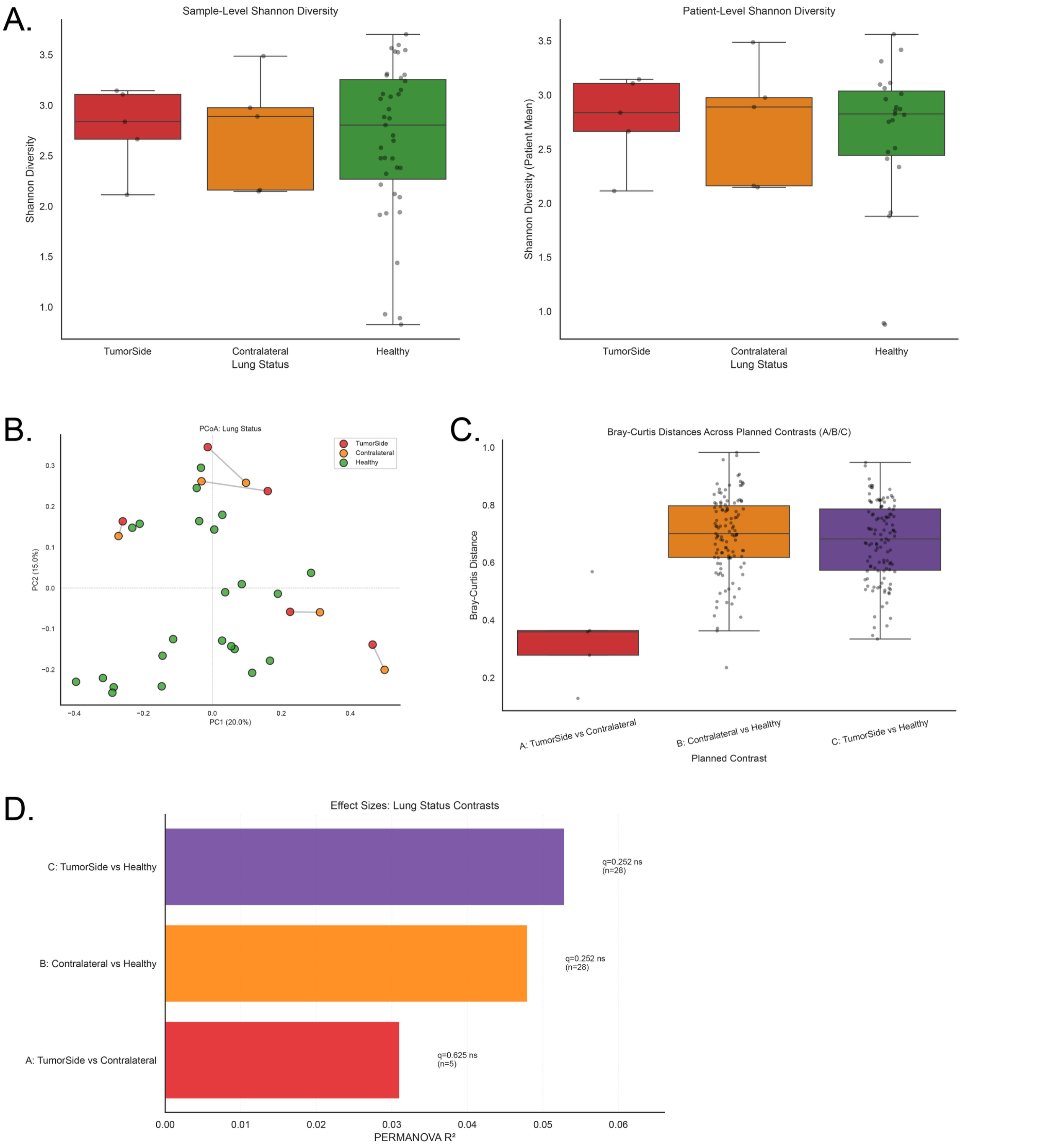


**Supplementary Figure S3. Lower-airway community structure by lung status (bronchial brush).** Comparison of bronchial brush communities among tumour-side, contralateral (non-tumour) and healthy-control lungs. (A) Shannon alpha diversity by lung status; (B) principal-coordinates analysis (PCoA) of Bray–Curtis dissimilarities; (C) distribution of pairwise Bray–Curtis distances for each contrast; (D) PERMANOVA $R^2$ for each contrast. No PERMANOVA centroid or alpha-diversity contrast reached the prespecified significance threshold: tumour-side versus contralateral (paired, n = 5 patients; PERMANOVA $R^2 = 0.058$, $p = 0.88$), contralateral versus healthy (n = 28; $R^2 = 0.046$, $p = 0.14$), and tumour-side versus healthy (n = 28; $R^2 = 0.035$, $p = 0.52$). Multivariate dispersion (PERMDISP) differed for contralateral versus healthy ($p = 0.047$) and was borderline for tumour-side versus healthy ($p = 0.050$); these p-values are unadjusted. Here n denotes participants contributing bronchial brush samples to each comparison (23 healthy controls plus the relevant cancer group); paired tumour-side and contralateral brushes were available for 5 of the 8 cancer participants. Contralateral samples appeared intermediate between tumour-side and healthy profiles, but the comparison was underpowered (see Power Analysis). Full statistics are provided in Supplementary Table S1 and Supplementary Dataset SD5.

A.

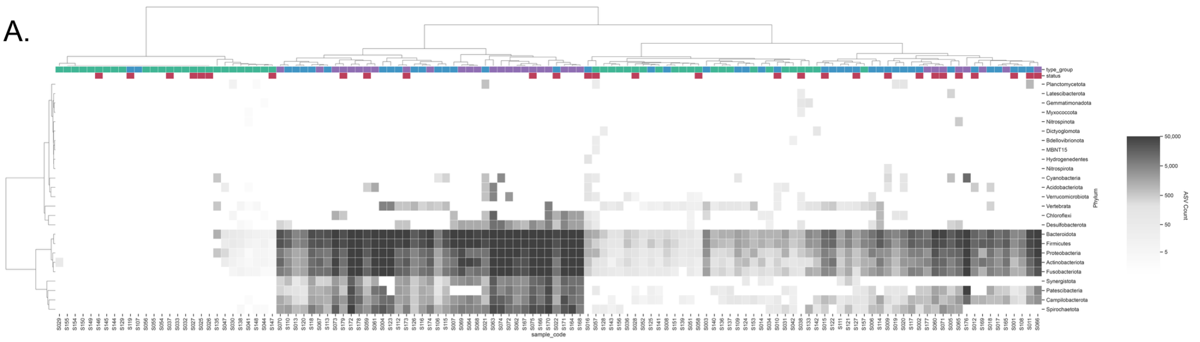


B.

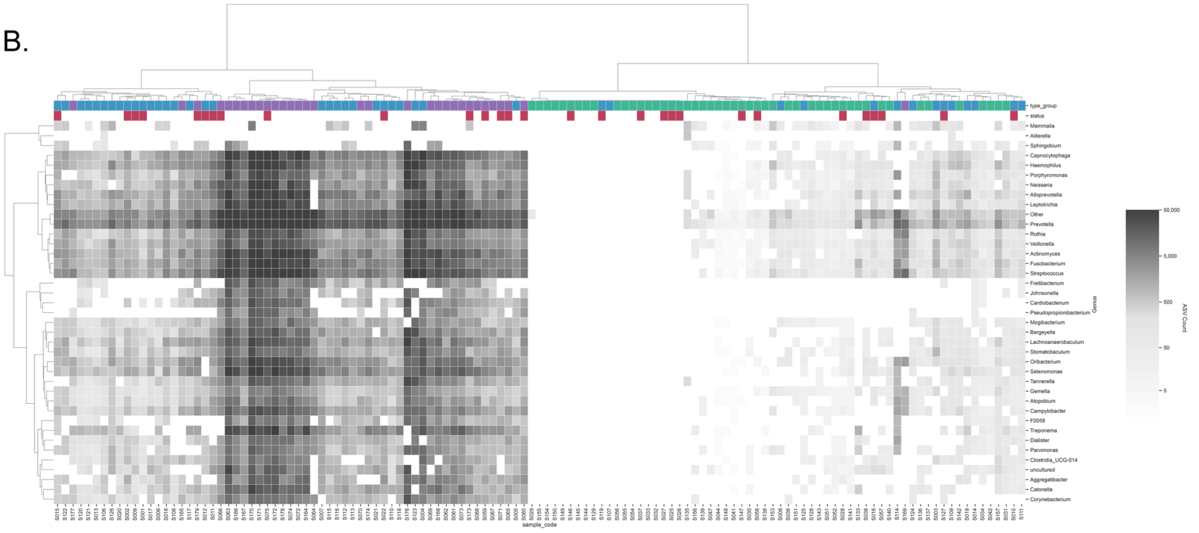


**Supplementary Figure S4. Hierarchical clustering of samples at additional taxonomic levels.** Hierarchical cluster analysis (Ward linkage on Euclidean distances) of sample relative-abundance profiles summarized at the (A) phylum and (B) genus levels, complementing the family-level clustermap shown in Figure 3B. Each column is one sample; the colored annotation bars above the heatmap denote sample type (Oral Rinse, purple; BAL, blue; Bronchial Brush, green) and cancer status (Non-Cancer, white; Cancer, red). At both taxonomic resolutions, samples resolved into two principal clusters (one dominated by low-read-count bronchial brush samples and the other by oral rinse and BAL samples) consistent with the family-level and UMAP results, and no cluster was enriched for cancer status.

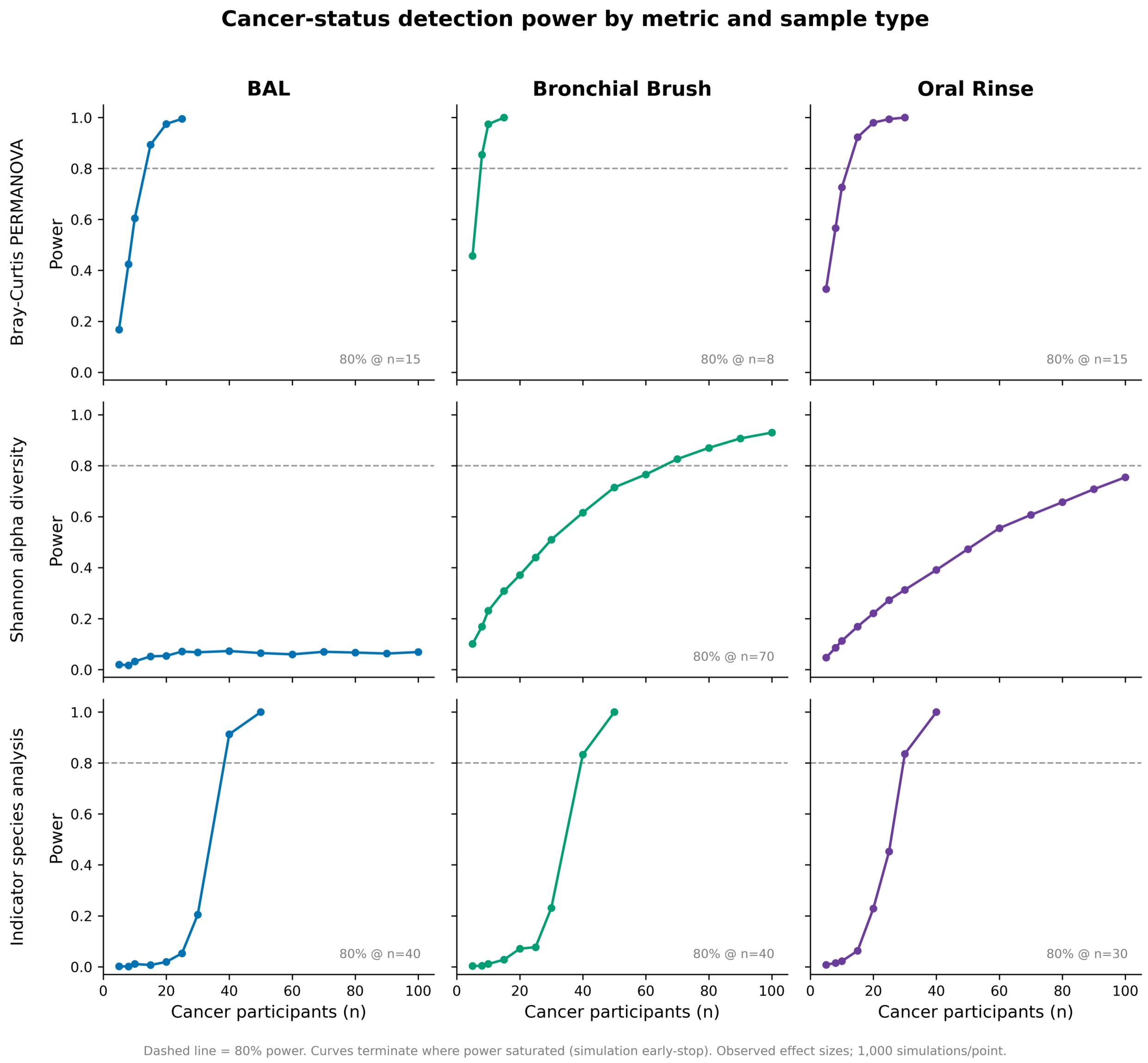


**Supplementary Figure S5. Cancer-status detection power by metric and sample type.** Simulation-based statistical power to detect cancer-versus-control differences as a function of cohort size (n = 5 to 100 cancer participants per sample type), estimated under the effect sizes observed in this cohort (1,000 simulations per point; α = 0.05). Rows correspond to three analytical approaches — Bray–Curtis PERMANOVA (community structure), Shannon alpha diversity, and indicator-species analysis (ISA; multipatt combination testing) — and columns to the three sample types (BAL, bronchial brush, oral rinse). The dashed line marks 80% power, and the cohort size at which each curve first reaches 80% is annotated. Curves terminate where power saturated (simulation halted once power reached ≥99.5%). Community-structure (Bray–Curtis) and indicator-species comparisons reached adequate power at moderate cohort sizes (n ≈ 8–15 and n ≈ 30–40, respectively), whereas Shannon alpha diversity remained underpowered across the tested range for BAL and oral rinse, reaching 80% only for bronchial brush (n ≈ 70) — consistent with the insensitivity of alpha diversity to the compositional differences that distinguish these communities.

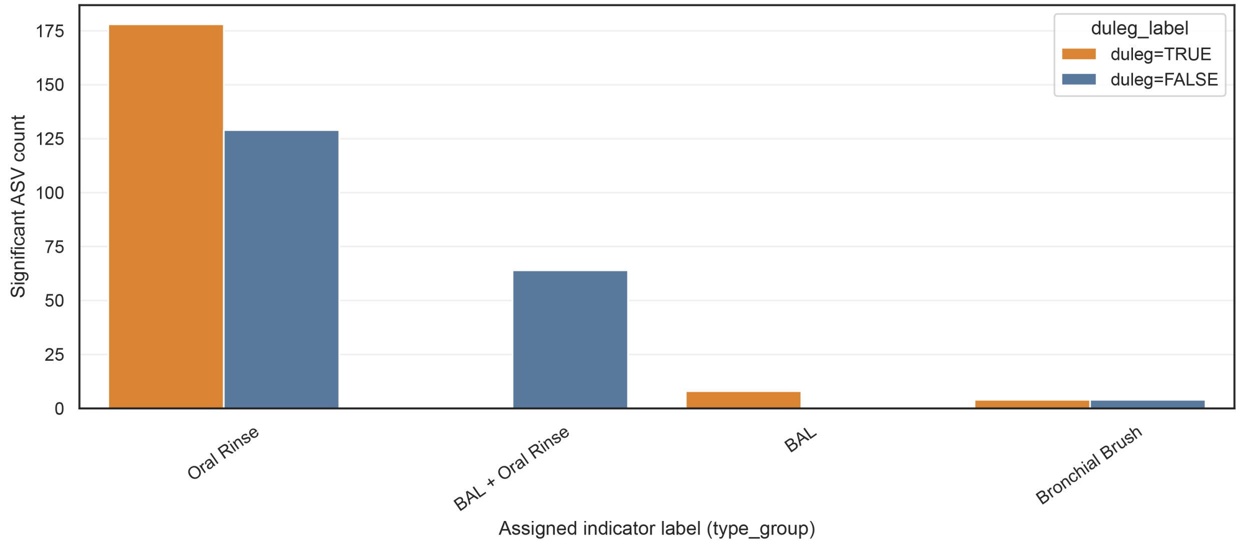


**Supplementary Figure S6. Distribution of sample-type indicator ASVs across habitat combinations.** Number of significant indicator ASVs (indicator stat ≥ 0.25, q < 0.05) assigned to each sample-type combination by multi-level indicator-species analysis. Of 197 significant indicators, 129 were associated with oral rinse alone, 64 jointly with BAL and oral rinse, and 4 with bronchial brush alone; no ASV was a joint BAL–bronchial brush indicator. Full indicator results are provided in Supplementary Dataset SD8a.

### 4. Supplementary Tables

| contrast | Comparison type | N patients | N samples | Permanova R2 | Permanova F | Permanova p | Permdisp F | Permdisp p | Alpha statistic | Alpha p | Alpha median group1 | Alpha median group2 | Permanova q | Alpha q |
|---|---|---|---|---|---|---|---|---|---|---|---|---|---|---|
| A TumorSide vs Contralateral | paired | 5 | 10 | 0.058 | 0.4923 | 0.875 | 0.0014 | 0.936 | 11 | 0.4185 | 2.9017 | 2.6418 | 0.875 | 0.6277 |
| B Contralateral vs Healthy | Between patient | 28 | 28 | 0.0459 | 1.2511 | 0.1381 | 3.9169 | 0.047 | 55 | 0.9045 | 2.6418 | 2.7933 | 0.4143 | 0.9045 |
| C TumorSide vs Healthy | Between patient | 28 | 28 | 0.0348 | 0.9383 | 0.5195 | 3.8846 | 0.05 | 41 | 0.3372 | 2.9017 | 2.7933 | 0.7792 | 0.6277 |

**Supplementary Table S1. Contralateral lung-status comparisons (bronchial brush).** Per-contrast PERMANOVA, PERMDISP, and alpha-diversity results among tumour-side, contralateral (non-tumour), and healthy-control lungs. No contrast reached significance.

| Df | SumOfSqs | R2 | F | Pr(>F) | term | model |
|---|---|---|---|---|---|---|
| 2 | 1.42 | 0.0574 | 2.62 | 0.001 | sample_type | sample_type_within_patient |
| 86 | 23.3 | 0.943 | NA | NA | Residual | sample_type_within_patient |
| 88 | 24.8 | 1 | NA | NA | Total | sample_type_within_patient |

| model | Df | SumOfSqs | R2 | F | p value | group1 | group2 | N patients | q-value |
|---|---|---|---|---|---|---|---|---|---|
| Sample type pairwise | 1 | 0.335 | 0.0214 | 1.1 | 0.001 | BAL | Bronchial Brush | 26 | 0.001 |
| Sample type pairwise | 1 | 0.632 | 0.0461 | 2.71 | 0.001 | BAL | Oral Rinse | 29 | 0.001 |
| Sample type pairwise | 1 | 0.991 | 0.0621 | 3.44 | 0.001 | Bronchial Brush | Oral Rinse | 27 | 0.001 |

| model | statistic | p_value | note |
|---|---|---|---|
| sample_type_within_patient | 16.7 | 0.001 | PERMDISP on Bray-Curtis (TSS) |
| case_status_patient_pooled | 20.1 | 0.001 | PERMDISP on Bray-Curtis (TSS) |
| case_status_BAL | 8.3 | 0.013 | PERMDISP on Bray-Curtis (TSS) |
| case_status_Bronchial Brush | 3.88 | 0.056 | PERMDISP on Bray-Curtis (TSS) |
| case_status_Oral Rinse | 6.81 | 0.012 | PERMDISP on Bray-Curtis (TSS) |

| group1 | group2 | n_paired_patients | wilcoxon_w | pval | qval | significant |
|---|---|---|---|---|---|---|
| Bronchial Brush | Oral Rinse | 27 | 13 | 1.3e-06 | 3.9e-06 | TRUE |
| Bronchial Brush | BAL | 26 | 45 | 4.7e-04 | 4.7e-04 | TRUE |
| Oral Rinse | BAL | 29 | 51 | 1.2e-04 | 1.9e-04 | TRUE |

**Supplementary Table S2. Diversity statistics by sample type.** (a) Patient-blocked Bray–Curtis PERMANOVA (global) and (b) pairwise comparisons; (c) PERMDISP test of multivariate dispersion homogeneity; (d) patient-aware pairwise Shannon alpha-diversity Wilcoxon tests. All p-values BH-FDR–corrected; PERMANOVA used 999 permutations (seed 42).

| Test | Sample type | n=5 | n=8 | n=10 | n=15 | n=20 | n=25 | n=30 | n=40 | n=50 | n=60 | n=70 | n=80 | n=90 | n=100 | Reached 80pct | n at 80pct | Max n simulated |
|---|---|---|---|---|---|---|---|---|---|---|---|---|---|---|---|---|---|---|
| Bray-Curtis PERMANOVA | BAL | 0.168 | 0.424 | 0.605 | 0.893 | 0.975 | 0.995 | | | | | | | | | yes | 15 | 25 |
| Bray-Curtis PERMANOVA | Bronchial Brush | 0.457 | 0.854 | 0.974 | 1 | | | | | | | | | | | yes | 8 | 15 |
| Bray-Curtis PERMANOVA | Oral Rinse | 0.327 | 0.566 | 0.726 | 0.923 | 0.98 | 0.994 | 1 | | | | | | | | yes | 15 | 30 |
| Indicator species (ISA) | BAL | 0.002 | 0.001 | 0.011 | 0.007 | 0.019 | 0.053 | 0.205 | 0.913 | 1 | | | | | | yes | 40 | 50 |
| Indicator species (ISA) | Bronchial Brush (no contralateral) | 0.003 | 0.004 | 0.011 | 0.028 | 0.071 | 0.077 | 0.231 | 0.833 | 1 | | | | | | yes | 40 | 50 |
| Indicator species (ISA) | Oral Rinse | 0.008 | 0.015 | 0.022 | 0.063 | 0.229 | 0.453 | 0.836 | 1 | | | | | | | yes | 30 | 40 |
| Shannon alpha diversity | BAL | 0.02 | 0.017 | 0.032 | 0.052 | 0.054 | 0.071 | 0.068 | 0.073 | 0.065 | 0.06 | 0.07 | 0.067 | 0.063 | 0.069 | no | not reached | 100 |
| Shannon alpha diversity | Bronchial Brush | 0.101 | 0.169 | 0.231 | 0.308 | 0.371 | 0.44 | 0.51 | 0.616 | 0.715 | 0.766 | 0.826 | 0.87 | 0.907 | 0.93 | yes | 70 | 100 |
| Shannon alpha diversity | Oral Rinse | 0.047 | 0.086 | 0.113 | 0.169 | 0.221 | 0.273 | 0.313 | 0.391 | 0.473 | 0.555 | 0.607 | 0.657 | 0.708 | 0.755 | no | not reached | 100 |
| Taxonomic abundance (Family) | BAL | 0.376 | 0.771 | 0.888 | 0.99 | 1 | | | | | | | | | | yes | 10 | 20 |
| Taxonomic abundance (Family) | Bronchial Brush | 0.262 | 0.428 | 0.615 | 0.94 | 0.994 | 1 | | | | | | | | | yes | 15 | 25 |
| Taxonomic abundance (Family) | Oral Rinse | 0.18 | 0.321 | 0.445 | 0.731 | 0.919 | 0.975 | 0.997 | | | | | | | | yes | 20 | 30 |
| Taxonomic abundance (Phylum) | BAL | 0.081 | 0.132 | 0.177 | 0.309 | 0.459 | 0.663 | 0.867 | 0.973 | 0.997 | | | | | | yes | 30 | 50 |
| Taxonomic abundance (Phylum) | Bronchial Brush | 0.443 | 0.635 | 0.819 | 0.986 | 0.998 | | | | | | | | | | yes | 10 | 20 |
| Taxonomic abundance (Phylum) | Oral Rinse | 0.215 | 0.311 | 0.397 | 0.574 | 0.711 | 0.793 | 0.876 | 0.975 | 0.999 | | | | | | yes | 30 | 50 |

**Supplementary Table S3. Statistical power for cancer-status comparisons.** Simulation-based power (1,000 simulations per point; observed effect sizes) for cancer-versus-control comparisons by test (Bray–Curtis PERMANOVA, Shannon alpha diversity, phylum- and family-level abundance, indicator-species analysis) and sample type, across cohort sizes from n=5 to n=100 cancer participants. Observed power should be interpreted as an upper bound owing to effect-size inflation in small cohorts. Blank cells indicate sample sizes not simulated; simulation halted once power reached ≥99.5%. Power was estimated from 1,000 simulations per cell under the effect sizes observed in this cohort.

| ASV_ID | Genus | Family | Indicator status | On Salter2014 | On Eisenhofer2019 |
|---|---|---|---|---|---|
| ASV1479 | Mammalia | Mammalia | Bronchial Brush | no | no |
| ASV1544 | Massilia | Oxalobacteraceae | | yes | yes |
| ASV168 | Bergeyella | Weeksellaceae | | not assessed | not assessed |
| ASV1907 | Acetomicrobium | Synergistaceae | | no | no |
| ASV2345 | Mammalia | Mammalia | | no | no |
| ASV325 | Sphingobium | Sphingomonadaceae | Bronchial Brush | yes | yes |
| ASV347 | Achromobacter | Alcaligenaceae | | no | yes |
| ASV45 | | Microbacteriaceae | | no | no |
| ASV587 | Oribacterium | Lachnospiraceae | | not assessed | not assessed |
| ASV651 | Acidovorax | Comamonadaceae | | yes | yes |
| ASV655 | Acinetobacter | Moraxellaceae | | yes | yes |
| ASV829 | Mammalia | Mammalia | Bronchial Brush | no | no |
| ASV873 | Sphingopyxis | Sphingomonadaceae | | yes | no |
| ASV888 | uncultured | Steroidobacteraceae | | no | no |
| ASV956 | Rhodoferax | Comamonadaceae | | no | no |
| ASV957 | Malikia | Comamonadaceae | | no | no |
| ASV973 | Mammalia | Mammalia | Bronchial Brush | no | no |

**Supplementary Table S4. Bronchial brush–module taxa cross-referenced against published contaminant lists.** The 17 members of the bronchial brush–associated co-occurrence network module, showing genus, family, indicator status, and whether each genus appears on the Salter et al. (2014) and/or Eisenhofer et al. (2019) reagent- and laboratory-contaminant lists. Sphingobium, Acidovorax, Acinetobacter, and Massilia appear on both lists; Sphingopyxis on Salter only; Achromobacter on Eisenhofer only; Rhodoferax, Malikia, and the uncultured Steroidobacteraceae indicator on neither.

### 5. Supplementary Datasets

**Supplementary Dataset SD1. ASVs removed by contamination filtering.**

The 37 ASVs removed from the analyzed set by the three-tier contamination filter, listed with the filter tier(s) responsible for each (decontam prevalence, within-sample-type frequency, and/or biological-plausibility screen for thermophilic environmental taxa), full taxonomy, and decontam p-values. (Zenodo, DOI: 10.5281/zenodo.22839013.)

**Supplementary Dataset SD2. Per-sample sequencing and quality-control summary.**

Read counts, observed ASV richness, sample type, cancer status, and participant ID for each of the 125 analyzed samples. (Zenodo, DOI: 10.5281/zenodo.22839013.)

**Supplementary Dataset SD3. Filtered ASV count table.**

ASV-by-sample count matrix for the 1,342 analyzed ASVs across 125 samples (the input to all downstream analyses). Available via Zenodo (DOI: 10.5281/zenodo.22839013).

**Supplementary Dataset SD4. ASV sample-type membership.**

Sample-type compartment assignment (which combination of oral rinse, BAL, and bronchial brush each ASV occurs in), total reads, and full taxonomy for all 1,342 ASVs. (Zenodo, DOI: 10.5281/zenodo.22839013)

**Supplementary Dataset SD5. Patient-level Bray–Curtis distance matrix for the contralateral lung-status analysis.**

Symmetric 33 × 33 matrix of patient-level Bray–Curtis dissimilarities between all bronchial brush samples included in the contralateral analysis, spanning healthy-control, contralateral (non-tumour) and tumour-side lungs. Sample labels denote participant ID and lung status (Healthy, Contralateral, or TumorSide). These distances underlie the lung-status contrasts summarized inSupplementary Table S1 and Supplementary Figure S3. (Zenodo, DOI: 10.5281/zenodo.22839013)

**Supplementary Dataset SD6. Taxonomic relative abundance by sample type.**

Mean relative abundance of every phylum and family in oral rinse, BAL, and bronchial brush samples. (Zenodo, DOI: 10.5281/zenodo.22839013)

**Supplementary Dataset SD7. Differential taxonomic abundance between sample types**.

(a) Friedman omnibus tests and (b) pairwise post-hoc comparisons across sample types at phylum and family level, with BH-FDR–corrected q-values. (Zenodo, DOI: 10.5281/zenodo.22839013)

**Supplementary Dataset SD8. Indicator-species analysis (complete results).**

Full multi-level indicator-species output for (a) sample-type and (b) cancer-status groupings, including indicator statistic, p- and q-values, and group assignment for every tested ASV. No ASV was a significant cancer-status indicator. (Zenodo, DOI: 10.5281/zenodo.22839013)

**Supplementary Dataset SD9. Co-occurrence network module membership.**

Module assignment, taxonomy, and centrality metrics (degree, betweenness, closeness, eigenvector) for all 278 nodes of the SPIEC-EASI network. (Zenodo, DOI: 10.5281/zenodo.22839013)

**Supplementary Dataset SD10. Representative ASV sequences**.

FASTA-formatted representative sequences for all 1,342 analyzed ASVs, with SILVA lineage in the sequence headers. ASV identifiers are internally consistent across SD1–SD10 but are not stable across pipeline runs; SD10 provides the mapping from identifier to sequence. (Zenodo, DOI: 10.5281/zenodo.22839013)
